# Wide sampling of natural diversity identifies novel molecular signatures of C_4_ photosynthesis

**DOI:** 10.1101/163097

**Authors:** Steven Kelly, Sarah Covshoff, Samart Wanchana, Vivek Thakur, W. Paul Quick, Yu Wang, Martha Ludwig, Richard Bruskiewich, Alisdair R. Fernie, Rowan F. Sage, Zhijian Tian, Zixian Yan, Jun Wang, Yong Zhang, Xin-Guang Zhu, Gane Ka-Shu Wong, Julian M. Hibberd

## Abstract

Much of biology is associated with convergent traits, and it is challenging to determine the extent to which underlying molecular mechanisms are shared across phylogeny. By analyzing plants representing eighteen independent origins of C_4_ photosynthesis, we quantified the extent to which this convergent trait utilises identical molecular mechanisms. We demonstrate that biochemical changes that characterise C_4_ species are recovered by this process, and expand the paradigm by four metabolic pathways not previously associated with C_4_ photosynthesis. Furthermore, we show that expression of many genes that distinguish C_3_ and C_4_ species respond to low CO_2_, providing molecular evidence that reduction in atmospheric CO_2_ was a driver for C_4_ evolution. Thus the origin and architecture of complex traits can be derived from transcriptome comparisons across natural diversity.

## Main text

The evolution of complex traits has produced great diversity in form and function across the living world. A large number of similar complex traits have evolved independently in multiple disparate lineages indicating that common responses to environmental selection can result in convergent phenotypes^1,2^. The C_4_ photosynthetic pathway, with at least 65 independent origins distributed across the angiosperms^3^, is considered one of the most remarkable examples of evolutionary convergence in eukaryotes. Thus the C_4_ pathway represents an attractive trait with which to determine whether phylogenetically diverse species can be examined to discover the shared molecular basis of complex convergent phenotypes.

Using a comparative approach, we analyzed gene expression of 30 C_4_ and 17 C_3_ species representing 18 independent evolutionary origins of C_4_ photosynthesis (Fig. 1A,B). This set of species includes representatives from all seven orders within the eudicotyledons known to have evolved C_4_ photosynthesis (Fig. 1a, Supplemental File 1)^4^. This sampling expands upon previous transcriptome studies with C_3_ and C_4_ eudicot plants in *Cleomaceae, Asteraceae* and *Portulaceae*^1^–^3^. RNA was isolated from leaves of all species, sequenced and subjected to *de novo* transcriptome assembly. Collectively these samples comprise 850 million reads that were assembled into 1.5 million contigs of which 1.1 million were assigned to orthogroups (Fig. 1B, Supplemental File 1) using a machine learning approach^5^. Analysis of the correlation in mRNA abundance estimates between species revealed that species did not cluster according to their photosynthetic type but rather according to phylogenetic relationship (Supplemental File 2). That is, C_4_ species of *Flaveria* are more similar to C_3_ *Flaveria* than to C_4_ species from other genera, and thus variation in gene expression underlying phenotypic convergence is not the primary determinant of differences in mRNA abundance between C_3_ and C_4_ species.

**Fig. 1:**
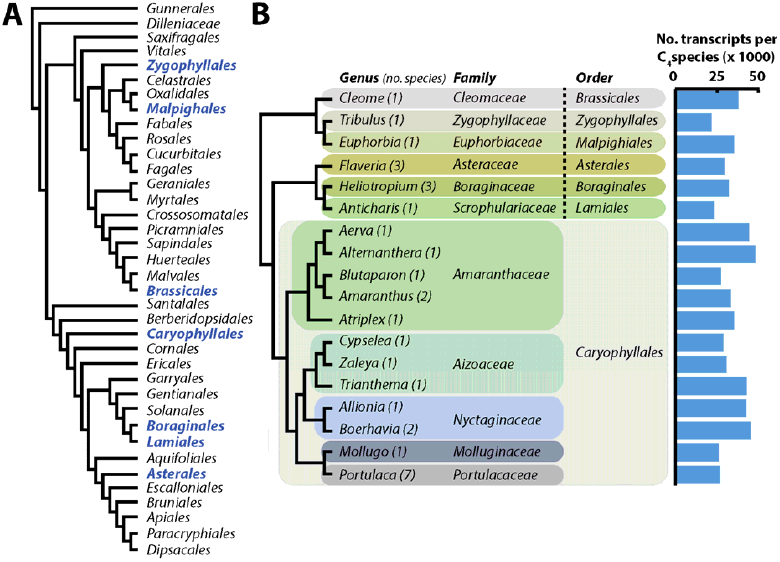
Within the eudicotyledons C_4_ photosynthesis has evolved in eleven families from 7 different orders. (A) Phylogenetic tree showing the seven orders containing C_4_ species (blue). (B) Phylogenetic tree showing the relationship between the eighteen genera (eleven families) encompassing thirty C_4_ species sampled in this study. Numbers after the genus name indicate the number of C_4_ species sampled. The mean number of *de novo* transcripts per species is indicated for each genus.

Comparison of the transcriptomes from C_3_ and C_4_ species identified 149 genes that showed altered transcript abundance between C_3_ and C_4_ species in all 18 lineages: 113 that were more abundant and 36 less abundant in all C_4_ species (Table 1, Supplemental File 3, Supplemental File 4). This set includes many genes encoding components of C_4_ photosynthesis that are known to change during evolution of the C_4_ pathway (Fig. 2A, Supplemental File 5, and Supplemental File 6). Four transcription factors were more abundant in all C_4_ species (*PAT1, ZML2, SHR* and a *bHLH* transcription factor of unknown function). Both *PAT1* and *ZML2* act to induce the expression of genes encoding photosynthesis proteins downstream of phytochrome and cryptochrome signalling respectively^6,7^ while *SHR* is a validated regulator of C_4_ Kranz anatomy in *Zea mays*^8^-^10^. Thus three of the four transcription factors have previously been identified as playing a role in the regulation of photosynthesis gene expression or leaf anatomy, both of which are altered during the evolution of C_4_ photosynthesis. The uncharacterised bHLH domain transcription factor has no known functional role, but has previously been described as being upregulated in the bundle sheath (BS) cells of the C_3_ plant *Arabidopsis thaliana*^11^. It is therefore possible that this bHLH transcription factor plays an ancestral role in the BS of C_3_ species that has become enhanced in all C_4_ lineages. Wide sampling of natural diversity therefore indicates that there is convergence in the recruitment of key regulators of gene expression in independent lineages of C_4_ species.

**Table 1:** Summary of functional categories of genes differentially expressed between C_4_ and C_3_ leaves.

| Category | Cell | Chloroplast | Mitochondrion |
| --- | --- | --- | --- |
| Metabolic components | 15 (8) | 29 (10) | 6 (4) |
| Signalling components | 18 (2) | 7 (1) | 0 (0) |
| Transporters | 4 (1) | 9 (1) | 2 (0) |
| Transcription regulators | 4 (1) | 0 (2) | 0 (0) |
| Post transcription regulators | 3 (1) | 0 (0) | 0 (0) |
| Other | 3 (0) | 2 (2) | 0 (0) |
| Unknown | 9 (3) | 1 (0) | 1 (0) |

n = 113, n = 36 No. up-regulated (No. down-regulated)

**Fig. 2:**
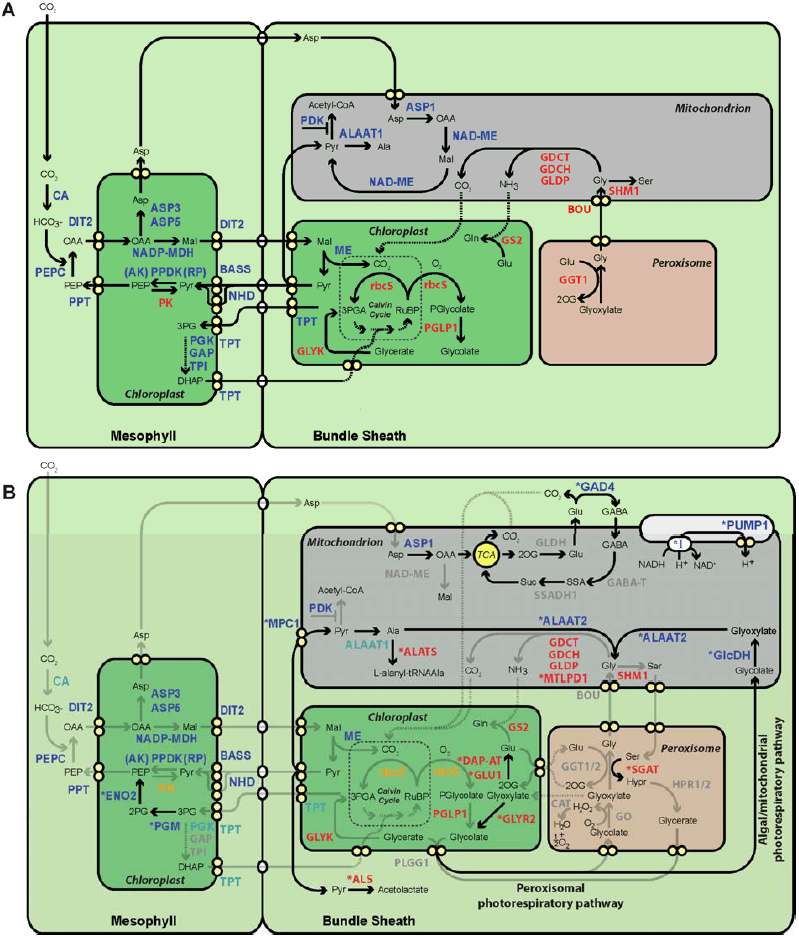
Schematics indicating proteins associated with C_4_ photosynthesis before (A) and after (B) this study. (A) Enzymes and transporters previously known to be upregulated (blue) and down-regulated (red) in C_4_ compared with C_3_ leaves are depicted according subcellular location and whether they are preferentially expressed in either mesophyll of bundle sheath cells. (B) After sequencing 30 C_4_ and 17 C_3_ species spanning the seven eudicotyledon orders known to have evolved C_4_ photosynthesis an additional 16 genes consistently up (blue) or down regulated (red) in C_4_ compared with C_3_ leaves were identified. Light blue and orange indicate transcripts for genes that are significantly upregulated and downregulated in all C_4_ species when compared to all C_3_, but they fail to achieve significance with computational occlusion and resampling. For abbreviations of gene names see Supplemental File 10. TCA, tricarboxylic cycle.

Transcripts encoding 16 proteins comprising four metabolic pathways that have not previously been associated with C_4_ photosynthesis were detected as differentially abundant between C_4_ and C_3_ species (Fig. 2B, Supplemental File 7). These pathways described below encompass: a novel carbon concentrating pathway involving the GABA shunt; metabolism associated with regeneration of phospho*enol*pyruvate (PEP), the primary CO_2_ acceptor in the C_4_ pathway; modifications to pyruvate metabolism that prevent diversion of pyruvate from the C_4_ cycle into non-photosynthetic pathways such as lipid and branched amino acid biosynthesis; and a photorespiratory pathway previously associated with chlorophyte algae (Supplemental File 7).

The abundance of transcripts encoding a key component of the γ-aminobutyric acid (GABA) shunt was increased in all C_4_ compared with C_3_ species. In the conventional model for NAD-ME type C_4_ photosynthesis, aspartate synthesised in mesophyll (M) cells is shuttled to mitochondria in the BS where it is transaminated to oxaloacetate by aspartate amino transferase (ASP1), reduced to malate by NAD-dependent malate dehydrogenase (NAD-MDH), and then decarboxylated to pyruvate which can then return to the M (Fig. 2a). Although *ASP1* transcripts were more abundant in all C_4_ species that we studied, this was not the case for the later steps in the NAD-ME pathway (Supplemental File 7). Instead, we propose that oxaloacetate is used to feed the tricarboxylic acid (TCA) cycle in BS cells. Here, 2-oxoglutarate synthesised by the TCA cycle can be converted to glutamate and decarboxylated by glutamate decarboxylase (GAD4) to GABA, resulting in release of CO_2_ and return of carbon skeletons as succinate to the TCA cycle (Fig. 2, Supplemental File 5, Supplemental File 7). This proposed pathway provides both a novel mechanism to transfer CO_2_ to the BS using CO_2_ that was fixed by phospho*enol*pyruvate carboxylase (PEPC) in the M, and a source of ATP in the BS by using NADH generated by the running the TCA cycle for oxidative phosphorylation (Fig. 2B, Supplemental File 5, Supplemental File 6). Two orthogonal approaches provide evidence that this cycle functions to concentrate CO_2_ in the C_4_ BS. First, computational modelling revealed that the additional ATP this pathway provides to BS cells resulted in an increased CO_2_ assimilation rate irrespective of C_4_ subtype under low light conditions (Supplemental File 8). Moreover, when PEPCK is used for decarboxylation this increase in CO_2_ assimilation rate was maintained under high light (Supplemental File 8). Second, biochemical evidence for this pathway has in fact been reported previously - after ^14^C labelled glutamate was fed to isolated BS strands in *Zea mays*^12^, radiolabel is rapidly released and redistributed to other metabolites in a manner that is most parsimoniously explained by glutamate decarboxylase mediated decarboxylation followed by re-fixation of labelled CO^2^ by RuBisCO. Thus, sampling the natural diversity of C4 species uncovered an adjunct CO_2_ concentrating pathway that is supported by biochemical data and a metabolic model of C_4_ photosynthesis.

To maintain flux through the C_4_ pathway, PEP supply is critical as it is the entry point of the cycle. Transcripts encoding a chloroplastic phosphoglucomutase (PGM) and an enolase (ENO) were more abundant in all C_4_ compared with C_3_ species (Fig. 2B, Supplemental File 7). These proteins facilitate conversion of Calvin-Benson cycle intermediates to PEP (Fig. 2B), providing an additional route for transfer of photo-assimilated carbon to the M, and consequently regeneration of the initial carbon acceptor. Transcripts encoding pyruvate kinase (PK), which catalyzes the reverse reaction, were less abundant in C_4_ compared with C_3_ species (Fig. 2B). A reduction in the amount of the cognate protein would limit futile cycling between PEP and pyruvate during C_4_ photosynthesis. The third pathway detected in our analysis indicates pyruvate metabolism has been modified to prevent diversion of pyruvate from C_4_ photosynthesis into non-photosynthetic pathways such as lipid and branched amino acid biosynthesis. Transcripts encoding pyruvate dehydrogenase kinase (PDK) were more abundant (Fig. 2B), and aceto-lactate synthase (ALS) less abundant, in all C_4_ compared with C_3_ leaves (Supplemental File 7). As PDK deactivates pyruvate dehydrogenase by phosphorylation and ALS channels pyruvate into the synthesis of branched-chain amino acids, these alterations would support the core C_4_ cycle by reducing loss of pyruvate from photosynthetic pools.

The fourth pathway that we propose is modified in all C_4_ compared with C_3_ species has been previously associated with algae rather than land plants. Chloroplasts of chlorophyte algae enclose their RuBisCO in structures called pyrenoids. These structures facilitate an increased CO_2_ concentration around RuBisCO resulting in reduced photorespiration^13^. These algae also lack the peroxisome-based photorespiratory pathway that evolved in the common ancestor of embryophytes and charophyte algae^14^. Although the ancestral chlorophyte photorespiratory pathway involving glycolate dehydrogenase (GlcDH) and an alanine:glyoxylate amino transferase (ALAAT2) is still active in C_3_ plants, flux of glycolate through this pathway is low compared with the peroxisome-based pathway^15^. Our analysis indicates that in all 18 C_4_ lineages there is a concerted increase in abundance of transcripts encoding key components of the chlorophyte algal pathway, specifically GlcDH and ALAAT2 (Fig. 2B, Supplemental File 7). Two scenarios may explain this change during C_4_ evolution. First, the chlorophyte photorespiratory pathway plays a role in C_4_ photosynthesis. Second, GlcDH plays a role in converting some dihydroxyacetone phosphate (DHAP) produced in the Calvin-Benson cycle of BS cells to pyruvate *via* the methylglyoxal pathway enabling the use of some DHAP to maintain the C_4_ cycle (Supplemental File 9). In this latter scenario, ALAAT2 would still process photorespiratory glyoxylate that had been produced in the peroxisome by glycolate oxidase. Both proposed scenarios require a source of NAD^+^ and in this context it is noteworthy that both transcripts encoding Complex I of the respiratory electron transfer chain and the plant uncoupling mitochondrial protein 1 (PUMP1) were upregulated in C_4_ species (Fig. 2B). Together, these would increase regeneration of NAD^+^ and de-couple some proton flux through Complex I from ATP synthesis. Moreover, this increase in NAD^+^ would also support photorespiratory glycine decarboxylase and utilise NADH from the TCA cycle (Fig. 2B).

We also evaluated whether transcriptome sampling across a deep phylogeny could be used to clarify selective forces promoting C_4_ evolution. Low atmospheric CO_2_ has been proposed to be a key driver of C_4_ evolution^16^, and analysis of *A. thaliana* identified genes responsive to low CO_2_ in plants^17^. Thirty-one of the 113 genes that were more abundant in all C_4_ species sampled showed increased expression in *A. thaliana* grown under low CO_2_ (Fig. 3, Supplemental File 10). As the probability of such an overlap is low (*p* = 4 × 10^-9^), these data indicate that there is a significant association between genes that are expressed highly in C_4_ species and those that are more abundant in C_3_ *A. thaliana* grown under low CO_2_. There is also a significant association between genes that are more abundant under low CO_2_ and those that are less abundant in all C_4_ plants (*p* = 1 × 10 ^-5^, Fig. 3). However, nine of these twelve genes encode components of the photorespiratory pathway and the remaining three are unknown proteins predicted to localize to the chloroplast and are thus implicated in photorespiratory processes (Supplemental File 11). These results are consistent with the hypothesis that low atmospheric CO_2_ concentration induced changes in gene expression that facilitated C_4_ evolution. The molecular mechanisms that underpin this response remain to be identified. In the future it will be informative to investigate the extent to which other ecological drivers such as heat, drought and salinity alter gene expression and potentially target genes that are then recruited into this complex trait. In addition, the data also reveal that a set of genes that are more abundant in C_4_ species are preferentially expressed in the BS cells of C_3_ species and thus indicate that neofunctionalisation of BS cells utilized pathways already present in this cell type (Supplemental File 5, Supplemental File 12). Thus these data also provide molecular support for the hypothesis that expansion and specialization of the C_3_ BS is an early and key step in the evolution of the C_4_ phenotype^19,18^.

**Fig. 3:**
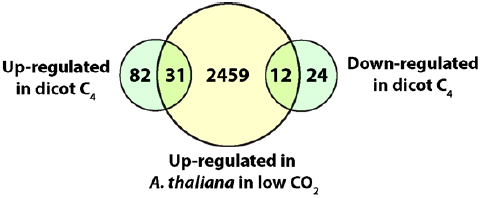
Overlap between genes that are upregulated in response to low CO_2_ in *Arabidopsis thaliana* and that are up or downregulated in C_4_ species.

Finally, the data identify four metabolic pathways previously unknown to be important for C_4_ function, and identify a role for the GABA shunt pathway in concentrating CO_2_ and generating ATP in the BS of all C_4_ species. The re-emergence in plants of the peroxisome-based photorespiratory pathway from algae is to our knowledge, the first documented example of an evolutionary reversion being a key component of the advent of complexity and convergence in eukaryotic biology. We envisage that this approach of comparative transcriptome sampling of non-model species will now be used to provide insight into molecular signatures associated with complex traits across the tree of life. The fact that we recapitulated previous knowledge of C_4_ photosynthesis, but also significantly extended the functional model of C_4_ metabolism, implies there is much more to be discovered about this pathway.

## Acknowledgements

This work was funded by a Bill and Melinda Gates Foundation and a Department for International Development award to IRRI and sub-awards to SK, XZ, RFS and JMH. The 1000 Plants (1KP) initiative, led by GKSW, is funded by the Alberta Ministry of Advanced Education, Alberta Innovates Technology Futures (AITF), Innovates Centre of Research Excellence (iCORE), Musea Ventures, BGI-Shenzhen and China National Genebank (CNGB), and DSERC Discovery grants to RFS. SK is a Royal Society University Research Fellow. This work was supported by the European Union’s Horizon 2020 research and innovation programme under grant agreement no 637765.

## List of Supplementary Materials

**Supplemental File 1:** Summary of C_3_ and C_4_ species sampled in this study. Each species is classified by its photosynthetic pathway, its family and order. The number of sequenced reads, assembled contigs, annotated contigs, and assembly N50 for each species obtained are also listed.

**Supplemental File 2:** Phylogenetic position accounts for more variance in mRNA abundance than photosynthetic pathway. The heat map depicts the Spearman’s ranked correlation coefficient (ρ) between each species pair computed from global mRNA abundance estimates. The hierarchical cluster (tree to the left of the heat map) was computed directly from the correlation coefficients by converting these correlation coefficients to distance estimates. The distance estimate between two species A and B is evaluated as 1 - ρ. i.e. *d*(A,B) = 1 - ρ. A tree is then inferred from these distance estimates using the minimum evolution principle. Names of C_4_ species are shown in blue.

**Supplemental File 3:** Summary of genes showing differential expression between C_4_ and C_3_ leaves. The likelihood of differential expression is provided considering all species. Also provided is the proportion of resampling tests in which the gene was detected as consistently differentially regulated between C_3_ and C_4_ species. The mean and standard deviation of the expression estimate is provided for the C_3_ and C_4_ cohort as well as the number of samples that were identified as outliers and masked prior to differential expression testing. The expected count for each gene for each species is also provided.

**Supplemental File 4:** The subset of genes from Supplemental File 3 that received 100% support from computational occlusion and resampling.

**Supplemental File 5:** Additional information.

**Supplemental File 6:** Genes previously reported to be differentially expressed in C_4_ compared with C_3_ leaves.

**Supplemental File 8:** Four additional metabolic pathways identified in this study. In each case the relative expression is given for C_3_ (grey bars) and C_4_ (green bars) species. ^*^ indicates a likelihood of differential expression ≥ 0.95 and 100% support from computational occlusion and resampling. n.s. indicates a non-significant difference between the C_3_ and C_4_ expression levels. (A) GABA shunt. (B) Phospho*enol*pyruvate regeneration. (C) First step of branched chain amino acid biosynthesis. (D) Chlorophyte photorespiratory pathway.

**Supplemental File 8:** Modelling the addition of the GABA shunt to the C4 photosynthesis.

**Supplemental File 9:** Schematic illustrating an alternative hypothesis for the function of the glycolate dehydrogenase gene that is potentially a lactate dehydrogenase. This pathway would convert dihydroxyacetone phosphate to pyruvate *via* methylglyoxal.

**Supplemental File 10:** Genes that were up-regulated in all C_4_ species and also up-regulated in response to low atmospheric CO_2_ in the C_3_ plant *Arabidopsis thaliana*.

**Supplemental File 11**: Genes that were down-regulated in all C_4_ species and also up-regulated in response to low atmospheric CO_2_ in the C_3_ plant *Arabidopsis thaliana*.

**Supplemental file 12:** The overlap between the genes differentially regulated in all C_4_ species with other datasets. A) Comparison of transcripts upregulated in all C_4_ species with those preferentially expressed in *Arabidopsis thaliana* bundlesheath cells. B) Comparison of genes identified as differentially abundant in all C_4_ species compared to those identified as differentially abundant between C_3_ and C_4_ species of *Flaveria*. C) Analysis of the cell type specific expression in *Zea mays* and *Setaria italica* of the orthologues of the genes identified as up-regulated in all C_4_ species in this study.

**Supplemental File 13:** Genes that were up-regulated in all C_4_ species and also up-regulated in in bundle sheath cells of the C_3_ plant *Arabidopsis thaliana*.

**Supplemental File 14:** Full names and accession numbers for all genes shown in Fig. 2.

## Supplementary Methods

Detailed descriptions of the data sources and methods.

## Author contributions

SK, GKSW, RB, RFS, SC and JMH conceived the work. SC and RFS acquired the plant collection and mRNA, while SK, VT, SW, WPQ, XGZ, YW, GKSW, ML, RB, JW, YZ, ZY, ZT, ARF and RFS conducted the analysis. SK designed and developed the bioinformatic analyses. SK and JMH interpreted the data and wrote the paper.

